# Mycoplasmal endosymbionts of *Trichomonas vaginalis* are associated with reduced risk for *Chlamydia trachomatis* endometrial infection in asymptomatic, coinfected, women

**DOI:** 10.64898/2026.08.31.748291

**Authors:** Xuejun Sun, Kacy S. Yount, Nefer N. Batsuli, Camille Campbell, Sangmi Jeong, Tammy Tollison, Xinxia Peng, Toni Darville, Xiaojing Zheng, Catherine M. O’Connell

## Abstract

*Trichomonas vaginalis* is a protozoan parasite that causes trichomoniasis, the most common curable non-viral sexually transmitted infection, while *Chlamydia trachomatis* is a bacterial pathogen that can ascend to the upper genital tract and cause pelvic inflammatory disease, infertility, and ectopic pregnancy. *T. vaginalis* harbors bacterial endosymbionts, including *Candidatus Malacoplasma girerdii*, an obligate symbiont, and *Metamycoplasma hominis*, which can live freely or symbiotically. In a 16S rRNA sequencing study of the cervicovaginal microbiome of at-risk women, *Ca. M. girerdii* abundance was one of 13 features predicting the absence of coincident upper tract infection, despite no direct association between *T. vaginalis* infection and reduced chlamydial ascension. Investigating the relationship between these microorganisms further, we found that *T. vaginalis* vaginal abundance correlated positively with chlamydial burden in women whose infection was confined to the cervix, whereas a nonsignificant inverse relationship was observed in women with endometrial spread. Among participants with high chlamydial burden, *Ca. M. girerdii* was detected exclusively in women without endometrial infection. Both endosymbionts trended toward more frequent detection, and higher abundance, in coinfected women without endometrial spread, while *M. hominis* abundance correlated strongly with *T. vaginalis* burden in this group. These findings suggest that mycoplasmal endosymbionts of *T. vaginalis*, rather than *T. vaginalis* itself, are microbial factors limiting chlamydial ascension, and point to a three-way interaction between parasite, endosymbiont, and bacterial pathogen that shapes upper genital tract *C. trachomatis* infection risk.

## Introduction

*Trichomonas vaginalis*, a unicellular, flagellated protist, causes trichomoniasis, the most common, curable non-viral sexually transmitted infection (STI), with an estimated 156 million new cases worldwide among people aged 15-49 in 2020 (1). Infection is often asymptomatic, yet it can damage vaginal and cervical tissue (2), raising the risk of endometritis and pelvic inflammatory disease (PID) (3), adverse pregnancy outcomes (4), infertility (5, 6), HIV acquisition (7), and cervical cancer (8).

*T. vaginalis* expresses adherence factors and innate immune modulators that induce proinflammatory cytokines (reviewed by Mercer and Johnson (9)) while disrupting tight junctions, to increase paracellular permeability (10). The parasite adheres to genital epithelial cells, draws on host-derived nutrients, and forms symbiotic relationships with viruses and with endosymbiotic bacteria of the order Mycoplasmoidales (11). Two species are of particular interest: *Candidatus* Malacoplasma girerdii, an obligate endosymbiont found only in association with *T. vaginalis* (12, 13), and *Metamycoplasma hominis*, a facultative endosymbiont capable of living freely in the cervicovaginal microbiome (CVM) or symbiotically within the parasite (14). Both associations alter*T. vaginalis* biology in vitro, affecting parasite growth, metabolism, and cytoadhesion, and can enhance the parasite’s ability to induce host cell production of proinflammatory cytokines (15–17). *T. vaginalis* infection is highly associated with elevated vaginal pH and a dysbiotic CVM (10, 18), characterized by diverse anaerobes and the absence of a dominant *Lactobacillus* species, conditions that also favor acquisition of other bacterial sexually transmitted pathogens such as *Neisseria gonorrhoeae* and *Chlamydia trachomatis*.

Given this shared susceptibility, a natural question is whether *T. vaginalis* coinfection — and resident endosymbionts — shape the course of chlamydial infection itself. In a 16S rRNA gene sequencing study of the CVM in women enrolled in the TRAC cohort, a well-characterized population at high risk for *C. trachomatis* infection (19), thirteen informative amplicon sequence variants (ASVs) predicted presence or absence of endometrial chlamydial infection (AUC = 0.74). *C. trachomatis* ASV abundance emerged as the strongest single predictor of chlamydial spread to the upper genital tract (20), consistent with earlier work showing that TRAC participants with endometrial infection carried a higher mean cervical chlamydial burden than those with infection confined to the cervix (P = .02) (21). Unexpectedly, abundance of an ASV assigned to *Ca. M. girerdii* predicted absence of chlamydial spread to the upper genital tract, despite prior studies finding no association between *T. vaginalis* infection itself and reduced chlamydial spread, in either asymptomatic women or those with PID (3, 21). *T. vaginalis* alone appeared irrelevant to chlamydial ascension, yet one of its bacterial endosymbionts seemed protective. This apparent paradox motivated the present study. It also raised the question of whether another *T. vaginalis* endosymbiont, *M. hominis*, might have a similar protective association limiting chlamydial ascension, despite not appearing as one of the original predictive features. One possibility is that its facultative lifestyle, alternating between free-living and symbiotic states, diluted the predictive association.

In this study, we asked whether mycoplasmal endosymbiosis of *T. vaginalis* explains protection from chlamydial ascension, and, if true, was this effect also mediated by the more frequently detected *M. hominis*. We found that *T. vaginalis* presence or overall abundance was not associated with altered risk of ascending chlamydial infection, reaffirming the earlier result. However, among *T. vaginalis*-infected women, there was a trend toward fewer endometrial infections among women with detectable *Ca.* M. girerdii and/or *M. hominis* ASVs, and among participants with high chlamydial burden, the *Ca.* M. girerdii ASV was detected exclusively in women without endometrial spread.

Together, these findings point to mycoplasmal endosymbionts as microbial factors most likely shaping chlamydial ascension risk for women coinfected with *T. vaginalis*, raising the question of how these bacteria alter the vaginal environment to constrain *C. trachomatis* from spreading to the upper genital tract.

## Materials and Methods

### Ethics approval and consent to participate

The study protocol was approved by the Institutional Review Boards for human research of the University of Pittsburgh (#PRO10010159) and the University of North Carolina (#13-3074). All participants provided written informed consent at the time of enrollment.

### T Cell Response against Chlamydia (TRAC) Cohort (246 participants)

This cohort was composed of young (median age, 21 years; range, 18-35 years) cis-gender women meeting any one of the following eligibility criteria: clinical evidence of mucopurulent cervicitis; diagnosis of gonorrhea or chlamydia prior to treatment; or reported sexual contact with a male who received a diagnosis of gonorrhea, chlamydia, or nongonococcal urethritis (19). Exclusion criteria included acute PID, pregnancy, uterine procedure or miscarriage in the preceding 60 days, menopause, hysterectomy, antibiotic therapy in the preceding 14 days, and allergy to study medications. TRAC participants were recruited from the Allegheny County Health Department’s Sexually Transmitted Diseases Clinic, the University of Pittsburgh Medical Center Magee-Womens Hospital (MWH) Ambulatory Care Clinic, and the Reproductive Infectious Disease Research Unit at MWH in Pittsburgh, PA, and enrolled into a longitudinal study designed to investigate T cell responses important for protection from incident chlamydial infection. Participants provided informed consent at the time of enrollment and completed questionnaires regarding obstetric/gynecologic history, behavioral practices, sex exposure, contraceptive methods, and symptoms. Clinical, histological, and microbiological testing was performed, and blood, cervicovaginal swabs, and endometrial samples were obtained at enrollment, after which all participants received single dose antibiotics to treat gonorrhea (ceftriaxone, 250 mg intramuscularly) and chlamydia (azithromycin, 1 g orally). Participants were assessed for cervical and endometrial *C. trachomatis* and *N. gonorrhoeae* infection at enrollment using AC2 NAAT (Hologic, Marlborough, MA), with overall cervical infection rates of 68% and 8.5%, respectively. Cervical *M. genitalium* and vaginal *T. vaginalis* infections were determined using the corresponding Aptima NAATs (Hologic) with overall infection rates of 17% and 17%, respectively. Participants were classified as CT+ at enrollment if cervical *C. trachomatis* infection was detected by NAAT. Among CT+ participants, those with positive endometrial *C. trachomatis* NAAT result at enrollment were classified as Endo+, whereas those with *C. trachomatis* at the cervix only were classified as Endo−.

### 16S library preparation and sequencing

DNA was extracted from cervicovaginal swabs stored at -80°C (20). The V4 region of 16S rRNA genes were amplified using the Illumina 16S V4 primer set of 515F [GTGYCAGCMGCCGCGGTAA] and 806R [GGACTACNVGGGTWTCTAAT] (22, 23) and sequencing libraries prepared, sequenced and analyzed as previously described (20).

### ASV taxonomy

As previously reported (20), inferred ASVs were assigned taxonomy based on two approaches: (i) using the sintax function in USEARCH (24) with the RDP Classifier 16S trainset No. 18 raw training database (https://sourceforge.net/projects/rdp-classifier/files/RDP_Classifier_TrainingData/) and (ii) using speciateIT with vSpeciateDB (https://github.com/ravel-lab/speciateIT) (25). Table 1 details the ASVs detected in the TRAC 16S rRNA dataset specific to this analysis. These include ASVs assigned to *C. trachomatis* and *Mycoplasmoides genitalium* based on RDP. ASV sequence # ZOtu45 was initially assigned as *Malacoplasma microti* (26) but subsequent examination (20) determined that the assigned ASV shared 100% identity with uncultured *Mycoplasma sp*. clone Mnola (27), currently *Candidatus Malacoplasma girerdii* (NCBI:txid1318617) (28). Four ASVs were assigned *Metamycoplasma hominis* according to both reference databases. Subsequent BLAST searching indicated that all showed top hits to *M. hominis.* Confidence in their accurate taxonomic assignment was further increased after alignment with two reference sequences for *M. hominis,* accession # AF443616 and M24473. This indicated that ZOtu1232, ZOtu1994, and ZOtu1995 showed variation at the 3’ end consistent with previously published reports (29, 30). In contrast to *C. trachomatis* which carries two copies of the 16S locus, members of the Mycoplasmoidales frequently carry only a single copy (31).

**Table 1.** Taxonomic assignment of Mycoplasma-associated ASVs detected in 16S rRNA libraries of TRAC participants.

| ASV # | RDP DB-based taxonomy assignment* using USEARCH |  | Confidence level |  | vSpeciateDB-based taxonomy assignment using speciateIT |  | Posterior probability | Consistency check<sup>‡</sup> | Final classification |
| --- | --- | --- | --- | --- | --- | --- | --- | --- | --- |
|  | genus | species | genus | species | genus | species |  |  |  |
| ZOtu42 | Metamycoplasma | *Metamycoplasma hominis* | 1.0 | 1.0 | Metamycoplasma | *Metamycoplasma hominis* | 0.97 | YES | *Metamycoplasma hominis* |
| ZOtu45 | Malacoplasma | *Malacoplasma microti* | 0.8 | 0.4 | Campylobacter | *Campylobacter fetus* | 0.23 | NO | *Ca_Malacoplasma gireddii* |
| ZOtu1129 | Mycoplasmaoides | *Mycoplasmaoides genitalium* | 1.0 | 0.8 | Pyramidobacter | *Pyramidobacter pisolens* | 0.26 | NO | *Mycoplasmaoides genitalium* |
| ZOtu1232 | Metamycoplasma | *Metamycoplasma hominis* | 1.0 | 1.0 | Metamycoplasma | *Metamycoplasma hominis* | 0.66 | YES | *Metamycoplasma hominis* |
| ZOtu1994 | Metamycoplasma | *Metamycoplasma hominis* | 1.0 | 1.0 | Metamycoplasma | *Metamycoplasma hominis* | 0.71 | YES | *Metamycoplasma hominis* |
| ZOtu1995 | Metamycoplasma | *Metamycoplasma hominis* | 0.8 | 0.8 | Metamycoplasma | *Metamycoplasma hominis* | 0.54 | YES | *Metamycoplasma hominis* |
| ZOtu76 | Chlamydia | *Chlamydia trachomatis* | 1.0 | 1.0 | Chlamydia | *Chlamydia trachomatis* | 0.48 | YES | *Chlamydia trachomatis* |

\*All ASVs, except for ZOtu76, were assigned to phylum: Tenericutes, class: Mollicutes order: Mycoplasmaoidales
<sup>‡</sup>Discordant assignments were resolved after direct alignment of ASVs with published 16S rRNA gene sequences from the relevant species (13).

### Quantitative PCR of T. vaginalis and C. trachomatis burden in genital tract specimens

*T. vaginalis* loads were measured by quantitative PCR using residual genomic DNA remaining after preparation of 16S libraries as template, with the previously published BTUB3f (TCCAAAGGTTTCCGATACAGT) and BTUB_bkmt reverse (GTTGTGCCGGACATAATCATG) primers (32) that target the highly conserved beta-tubulin (*BTUB*) loci. *C. trachomatis* burden had been quantified from residual cervical diagnostic sample using primers targeting the 16S rRNA locus as previously described (21).

### Detection and quantitation of cytokines in genital tract specimens

Cytokines were quantified in cervical secretions collected at enrollment from TRAC participants as previously reported (33) using Milliplex Magnetic Bead Assay Kits (Millipore Sigma). Following batch correction by ComBat method, values above the upper limit of quantification (ULOQ) were set to the respective ULOQ, and values below the lower limit of quantification (LLOQ) were set to half the respective LLOQ. Participants with *N. gonorrhoeae* or *M. genitalium* infections were excluded.

#### Statistics

Mycoplasmal ASVs were analyzed as both binary detection variables and continuous abundance variables. For binary analyses, ASV detection was defined as the presence of at least one sequence read (count > 0). For abundance analyses, ASV counts were transformed as log2(count + 1). Counts for four ASVs assigned to *M. hominis* were summed to estimate total *M. hominis*-associated ASV abundance of individual participants. Participants were classified into four groups based on detection of *M. hominis* and *Ca. M. girerdii* ASVs: *M. hominis* only, *Ca. M. girerdii* only, codetection of *M. hominis* and *Ca. M. girerdii*, and detection of neither endosymbiont.

The association between the presence or absence of mycoplasmal ASVs and *T. vaginalis* diagnostic across comparison groups was assessed using Pearson’s chi-square test. Fisher’s exact test was used when more than 20% of the expected cell counts were less than 5. Continuous variables, including log2-transformed ASV abundance, log10-transformed *T. vaginalis* burden, and log10-transformed *C. trachomatis* burden, were compared between two groups using non-parametric Wilcoxon rank-sum tests. These analyses were applied to comparisons by *T. vaginalis* status, *C. trachomatis* status, endometrial infection status, and mycoplasmal ASV detection group, as shown in Figures 1–3 and Supplementary Figure 1.

**Figure 1.**
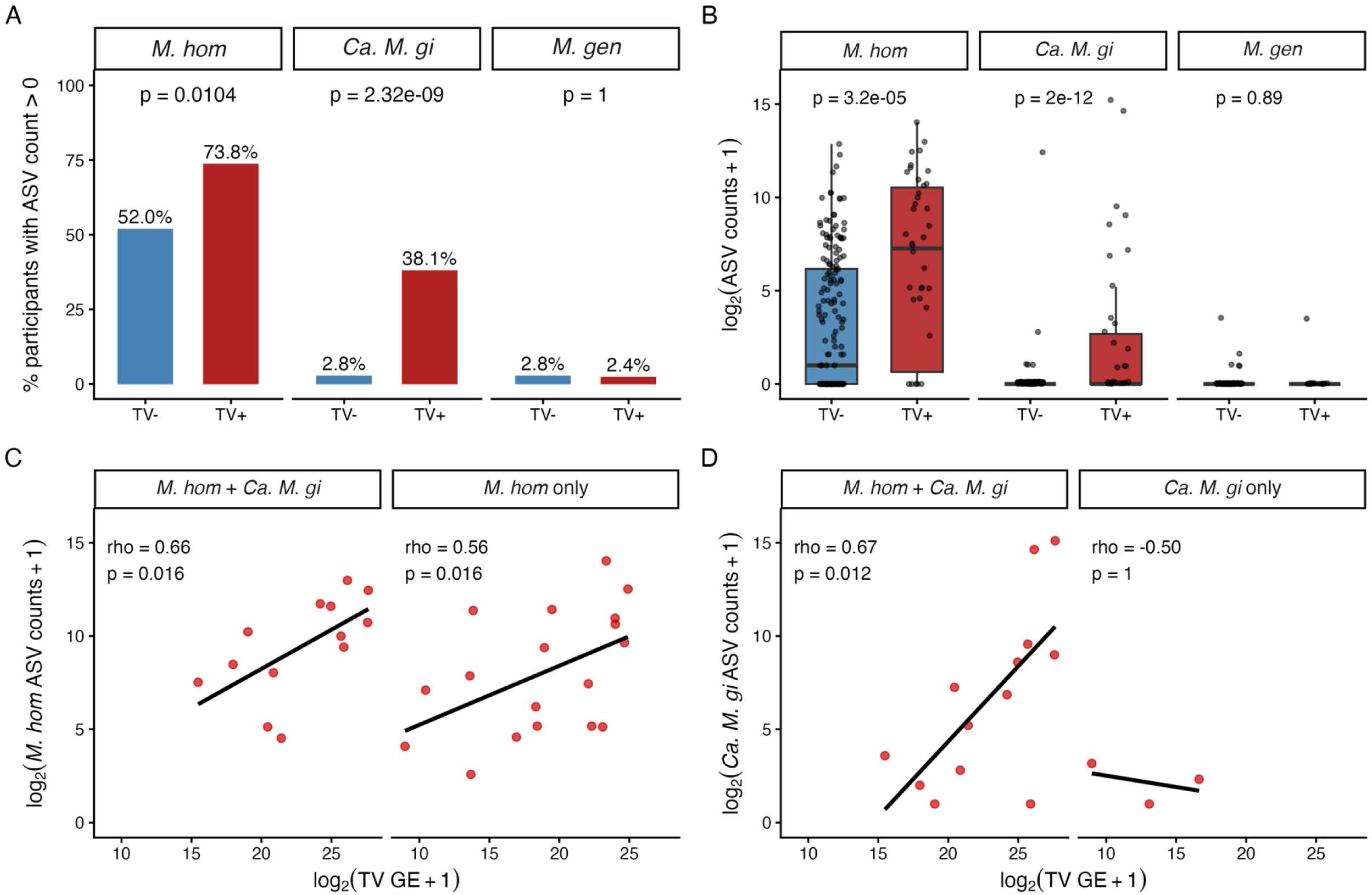
ASVs of Mycoplasmoidales capable of endosymbiosis with *T. vaginalis* (TV) are associated with TV infection status and abundance. (A) Percentage of participants with detectable ASV counts (>0) for *M. hominis* (*M. hom*), *Ca. M. girerdii* (*Ca. M. gi*), and *M. genitalium* (*M. gen*), stratified by TV status (TV− vs. TV+). P-values were obtained by chi-square test (Fisher’s exact test when expected counts were <5). (B) Distribution of log₂-transformed ASV counts (log₂[count + 1]) for *M. hom*, *Ca. M. gi*, and *M. gen*, stratified by TV status. Significance was assessed by Wilcoxon rank-sum test. (C) Association between log₂ *M. hom* ASV counts and log₂ TV burden (genome equivalents, GE) among TV-positive participants, stratified by participants carrying *M. hom* only versus *M. hom* and *Ca. M. gi*. Spearman correlation coefficients (ρ) and p-values are shown. (D) Association between log₂ *Ca. M. gi* ASV counts and log₂ TV burden (GE) among TV-positive participants, stratified by participants carrying *Ca. M. gi* only versus *M. hom* and *Ca. M. gi*. Spearman correlation coefficients (ρ) and p-values are shown.

**Figure 2.**
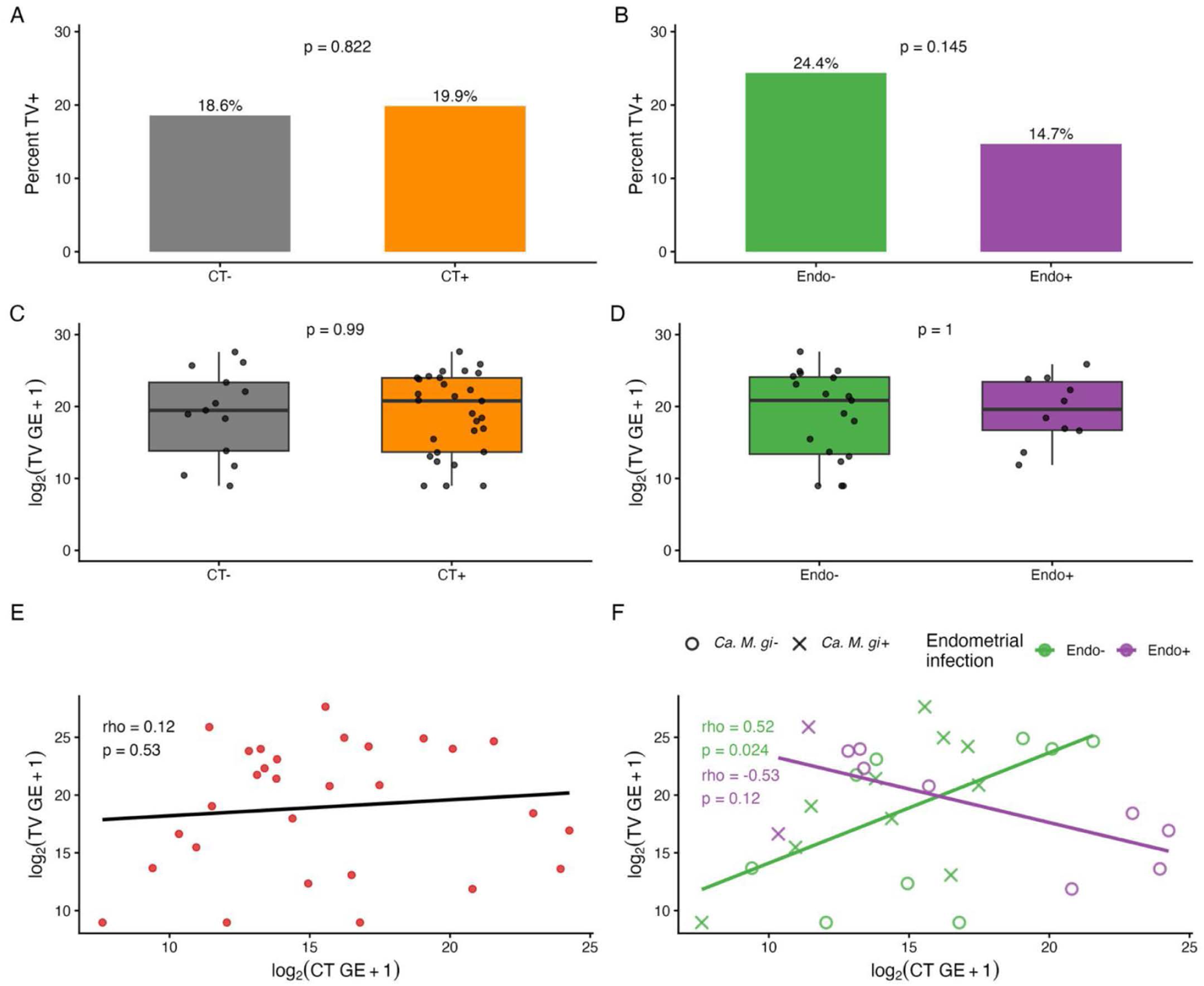
*T. vaginalis* (TV) presence and abundance are not associated with susceptibility to chlamydial infection or risk of endometrial spread, but chlamydial cervical burden correlates positively with trichomonad abundance among women without endometrial infection. Percentage of participants who were TV-positive, stratified by *C. trachomatis* (CT) cervical infection status (CT− vs. CT+) (A) and endometrial infection status (Endo− vs. Endo+) (B). Bars show the proportion TV-positive; P-values were obtained by chi-square or Fisher’s exact test. (C, D) Among TV-positive participants, TV burden (log₂[TV genome equivalents (GE) + 1]) compared between CT− and CT+ participants (C) and between Endo− and Endo+ participants (D). Boxplots display distributions; P-values were obtained by Wilcoxon rank-sum test. (E) Correlation between CT burden (log₂[CT GE + 1]) and TV burden (log₂[TV GE + 1]) among participants co-positive for CT and TV. Spearman’s ρ and P-value are shown. (F) Same as (E), stratified by chlamydial endometrial infection status (green = Endo−, purple = Endo+), with regression lines and Spearman correlations shown for each group.

**Figure 3.**
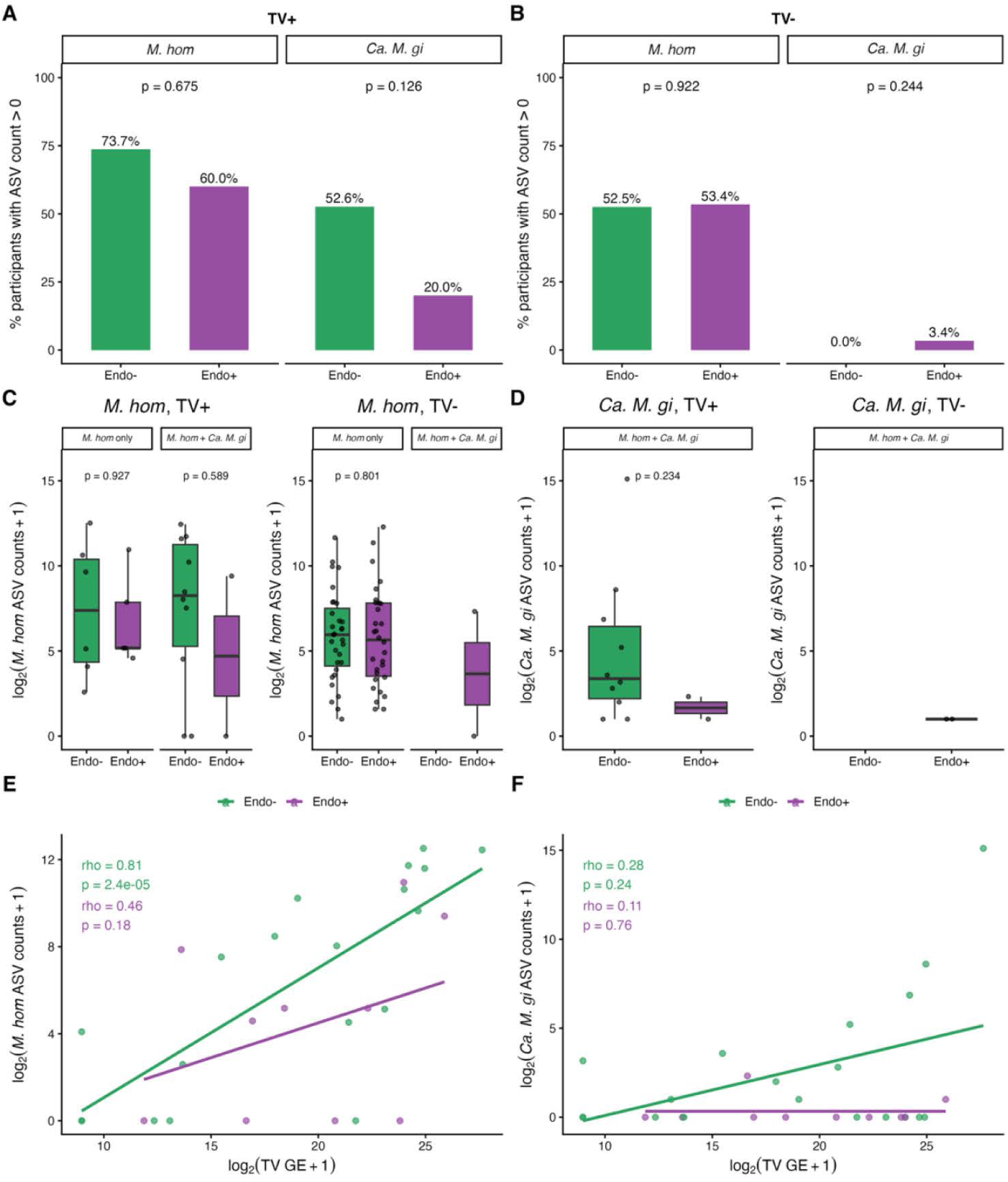
Frequency and abundance of endosymbiont-associated ASVs trend higher among Endo− TRAC participants coinfected with *T. vaginalis* (TV). (A) Percentage of *C. trachomatis* (CT)/TV-coinfected participants with detectable *M. hom* or *Ca. M. gi* ASV counts (>0), stratified by endometrial infection status (Endo− vs. Endo+). (B) Percentage of CT-infected, TV negative participants with detectable *M. hom* or *Ca. M. gi* ASV counts (>0), stratified by Endo− vs. Endo+. P-values in (A) and (B) were obtained by chi-square or Fisher’s exact test. (C, D) Among CT/TV-coinfected participants, abundance (log₂[count + 1]) of *M. hom* (C) and *Ca. M. gi* (D) ASVs, stratified by Endo− vs. Endo+. P-values were obtained by Wilcoxon rank-sum test. (E, F) Correlation between TV burden (log₂[TV genome equivalents + 1]) and ASV abundance for *M. hom* (E) and *Ca. M. gi* (F) among TV-positive participants, stratified by Endo− vs. Endo+. Lines show linear fits; Spearman’s ρ and P-values are shown for each group.

Associations between continuous microbial or pathogen burden measures were assessed using non-parametric Spearman’s rank correlation. These included associations between *T. vaginalis* burden and mycoplasmal ASV abundance, between *T. vaginalis* burden and *C. trachomatis* burden, and between *T. vaginalis* burden and *M. hominis* or *Ca. M. girerdii* abundance within strata defined by endometrial infection status. Scatterplots were generated with fitted linear regression lines to facilitate visualization of overall trends. Nevertheless, statistical associations were evaluated using Spearman’s rank correlation because microbial abundance and pathogen burden data were non-normally distributed. To evaluate whether the association between *Chlamydia trachomatis* (CT) burden and *Trichomonas vaginalis* (TV) burden differed by endometrial infection status, linear regression models were fitted with continuous TV burden as the outcome and CT burden, endometrial infection status, and the interaction term between CT burden and endometrial infection status as independent variables. A statistically significant interaction term indicated that the association between CT burden and TV burden differed according to endometrial infection status.

Cervical cytokine concentrations were compared between groups using non-parametric Kruskal-Wallis test followed by Dunn’s multiple comparisons test.

All tests were two-sided, and nominal *P*-values are reported. Statistical analyses were performed using R version 4.4.3 or GraphPad Prism v11.

#### Data access

The raw sequencing data are publicly available at NCBI’s Short Read Archive (SRA) under the BioProject accession number PRJNA1136868.

## Results

### ASVs assigned to *Ca. M. girerdii* and *M. hominis* are associated with *T. vaginalis* infection in TRAC participants

*T. vaginalis* infection at enrollment was detected by clinical diagnostic test, while *M. hominis* and *Ca. M. girerdii* carriage was inferred from ASVs in parallel 16S rRNA gene libraries (20). Of the 220 participants enrolled in the TRAC cohort, TV status was available for 219 (99.5%); 42 (19.2%) were TV+ and 177 (80.8%) were TV−. Endosymbiont carriage differed markedly by TV status (P = 3.03 × 10^-9^; Table 2): among TV+ participants, 18 (42.9%) carried MH alone, 3 (7.1%) carried MM alone, 13 (31.0%) carried both MH and MM, and 8 (19.0%) were negative for both endosymbionts, with complete endosymbiont data for all TV+ participants. Among TV− participants, MH alone was far more common (88, 49.7%) than either MM alone (1, 0.6%) or dual carriage (4, 2.3%), and nearly half (84, 47.5%) carried neither endosymbiont. Despite this striking difference in endosymbiont carriage, TV status itself was not associated with CT positivity (69.0% vs. 67.2% in TV+ vs. TV−; P = 0.822) or, among CT+ participants, with progression to endometrial infection (34.5% vs. 49.6%; P = 0.145).

**Table 2.**
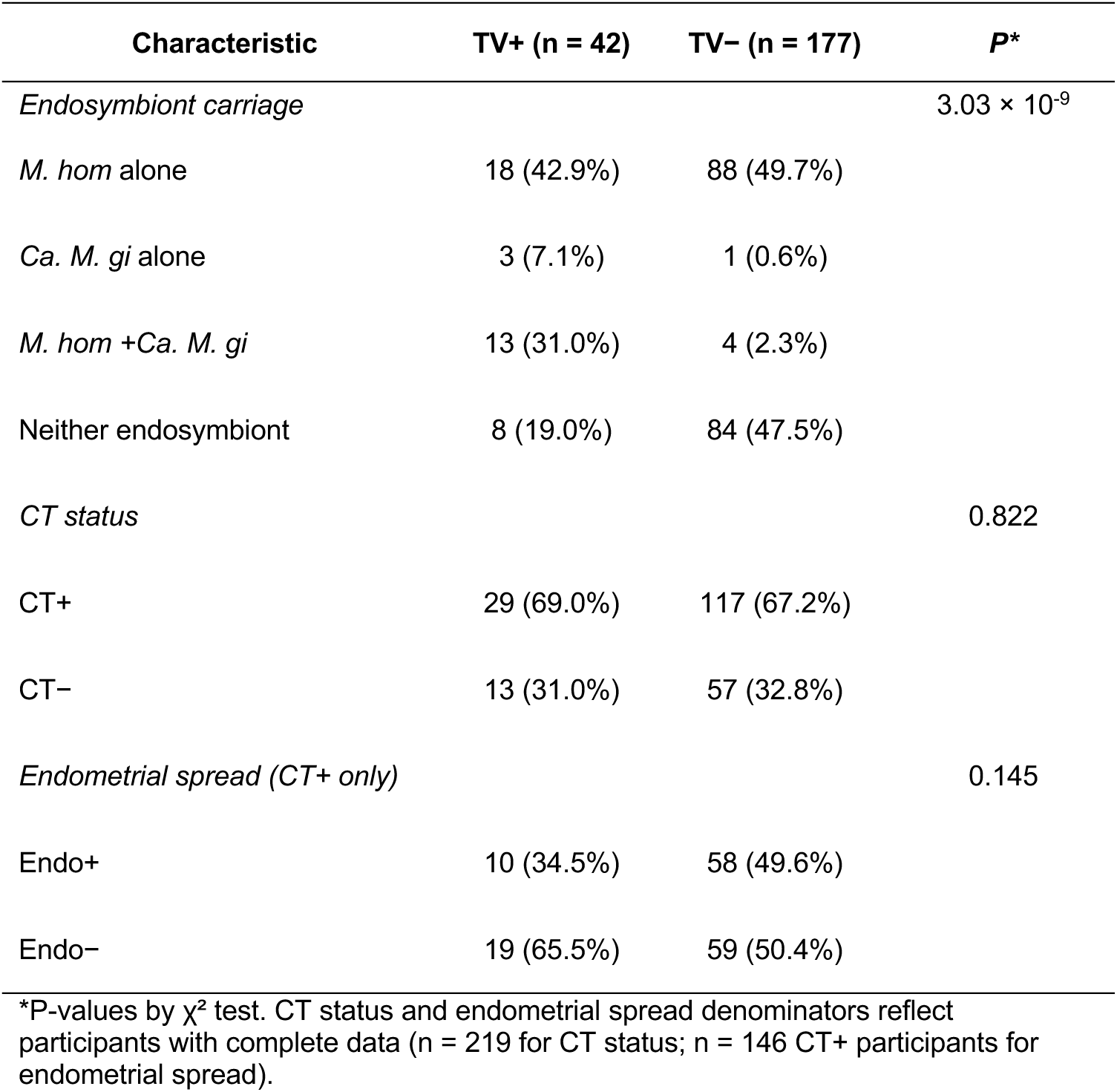
Participant TV, endosymbiont, and infection status.

Detection of the *Ca. M. girerdii*-assigned ASV was almost always associated with *T. vaginalis* infection (Fig. 1A, B), consistent with its lifestyle as a strict endosymbiont of the protist (12). Four ASVs assigned to *M. hominis* (Table 1) were summed to estimate overall *M. hominis* abundance per participant. *M. hominis*-assigned ASVs were detected in participants both with and without *T. vaginalis*, but more frequently (Fig. 1A) and at higher abundance (Fig. 1B) in those with trichomonads, consistent with its ability to live both freely and symbiotically (17). Forty-one TRAC participants were diagnosed with *M. genitalium*, a free-living sexually transmitted pathogen, by clinical assay(19, 34). The *M. genitalium*-assigned ASV was infrequently detected (N=6) and low in abundance (Fig. 1A, B), consistent with previous qPCR-based quantitation of *M. genitalium* cervical burden within the TRAC cohort (34) and displayed no evident association with *T. vaginalis* coinfection (N=11).

We next quantified vaginal *T. vaginalis* burden by qPCR, using residual template from 16S rRNA gene library synthesis. Endosymbiont presence promotes *T. vaginalis* growth in cell culture (16, 17) so we asked whether *T. vaginalis* abundance correlated with *M. hominis* and/or *Ca. M. girerdii* ASV counts. *M. hominis*-assigned ASVs were more frequently detected than the *Ca. M. girerdii*-assigned ASV, and coincident detection was common. *M. hominis*-assigned ASVs positively correlated with *T. vaginalis* abundance, both alone and in participants with coincident *Ca. M. girerdii* (Fig. 1C). Analyzing each of the four *M. hominis* ASVs individually (Supplementary Fig. 1)) gave similar results, indicating the correlation was not driven by a single ASV. Similarly, the *Ca. M. girerdii*-assigned ASV positively correlated with *T. vaginalis* burden in participants also carrying *M. hominis*, though no correlation was seen in the small *Ca. M. girerdii*-only subgroup (N=3) (Fig. 1D). Together, these data confirm that *Ca. M. girerdii* ASVs identified in the original 16S rRNA gene study reflected *T. vaginalis* infection, and that both *M. hominis*- and *Ca. M. girerdii-*assigned ASVs tracked with *T. vaginalis* abundance in the cervicovaginal environment of study participants.

### *T. vaginalis* abundance was positively associated with endosymbiont presence specifically in women with chlamydial infection confined to the cervix

*T. vaginalis* diagnosis frequency did not differ significantly between enrollees with and without *C. trachomatis*, indicating that *T. vaginalis* does not influence susceptibility to chlamydial infection (Fig. 2A). Consistent with our prior work (19) *T. vaginalis* diagnosis was not associated with increased risk of chlamydial ascension to the endometrium. Nevertheless, we observed a trend toward more frequent *T. vaginalis* coinfection among participants whose chlamydial infection was confined to the cervix (Endo–) compared with those with endometrial spread (Endo+) (Fig. 2B).

Among *T. vaginalis*-infected participants, *T. vaginalis* burden did not differ by chlamydial infection status (Fig. 2C), nor between Endo- and Endo+ participants (Fig. 2D), indicating that *C. trachomatis*, regardless of whether it was confined to the cervix or had spread to the endometrium, did not alter *T. vaginalis* abundance. Across all participants, *C. trachomatis* load and *T. vaginalis* abundance were not correlated overall (Fig. 2E) but this relationship differed by endometrial infection status (Fig. 2F); among Endo− participants, *C. trachomatis* load correlated positively with *T. vaginalis* abundance (rho = 0.52, p = 0.024), whereas among Endo+ participants the correlation was negative and nonsignificant (rho = −0.53, p = 0.12). A linear model of *T. vaginalis* abundance as a function of *C. trachomatis* load, endometrial infection status, and their interaction confirmed a significant interaction effect (p = 0.001), supporting a genuine difference in the *C. trachomatis*– *T. vaginalis* relationship between Endo− and Endo+ participants.

This pattern was notable given our previous 16S rRNA gene study, which identified the strict endosymbiont *Ca. M. girerdii* as a contributor to a cervicovaginal microbiome signature predicting the absence of coincident chlamydial endometrial infection (20) Chlamydial cervical burden is the strongest known correlate of endometrial spread (20, 21) so our observation that these Endo- participants tolerated high *C. trachomatis* and *T. vaginalis* burdens without spread suggested that some factor unique to this group buffered against chlamydial ascension despite high pathogen load. Among *T. vaginalis-*infected participants with high chlamydial burden (≥ 10^4^ GE), *Ca. M. girerdii* ASV was detected only in Endo− (7 of 13) and not in Endo+ (0 of 6) women (Fisher’s exact p = 0.044). In contrast, frequency of *Ca. M. girerdii* ASV detection was not different between Endo- (3 of 6) and Endo+ (2 of 4) among women with low chlamydial burden (< 10^4^ GE; Fisher’s exact p = 1.0). This suggested that, for these participants, coinfection with *T. vaginalis* harboring the *Ca. M. girerdii* endosymbiont might influence chlamydial upper genital tract ascension, despite the high chlamydial burden typically associated with increased spread. In further support, the *Ca. M. girerdii* ASV was detected 2.6-fold more often in Endo− than Endo+ groups among *T. vaginalis-*infected participants, though this difference did not reach significance (Fig. 3A, p = 0.126). ASVs representing the facultative endosymbiont *M. hominis* showed a similar but weaker trend toward more frequent detection in Endo− participants. In contrast, for chlamydia-infected participants without *T. vaginalis*, neither *Ca. M. girerdii* nor *M. hominis* ASVs differed in detection frequency between Endo− and Endo+ groups (Fig. 3B). As might be expected, the *Ca. M. girerdii* ASV was rarely detected at all, consistent with its strict dependency on *T. vaginalis* for replication (17).

Abundance of endosymbiont ASVs trended higher in coinfected Endo− participants (Fig. 3C–D). *M. hominis* abundance correlated strongly with *T. vaginalis* abundance among Endo− participants (rho = 0.81, p = 2.4e-05), but weakly in Endo+ participants (rho = 0.46, p = 0.18) (Fig. 3E) and this was not significant. This indicated that *M. hominis* had more consistently established an endosymbiotic relationship with the trichomonads infecting Endo− women. The *Ca. M. girerdii* ASV showed a similar but weaker pattern, a stronger positive correlation in Endo− than Endo+ participants (rho = 0.28, p = 0.24 vs. rho = 0.11, p = 0.76) (Fig. 3F), though neither correlation reached significance, likely reflecting the small number of participants harboring *Ca. M. girerdii* independent of *M. hominis*.

### Cervical cytokine levels were elevated by *T. vaginalis* and *C. trachomatis* infection, and IL-8 secretion was further increased by mycoplasmal endosymbiont detection

We quantified the proinflammatory cytokines MCP-1/CCL2, IL-6, IL-8, and TNFα in cervical secretions from TRAC participants stratified by *C. trachomatis* and *T. vaginalis* infection status and endosymbiont ASV detection (Fig. 4). Participants with *C. trachomatis* but without *T. vaginalis* (CT+TV−) had higher levels of MCP-1 (Fig. 4A), IL-6 (Fig. 4B), and TNFα (Fig. 4D) than participants with neither infection (CT−TV−). Among participants without *C. trachomatis* infection, those with *T. vaginalis* infection (CT−TV+) had similar MCP-1 levels (Fig. 4A) but higher IL-6 (Fig. 4B) and IL-8 (Fig. 4C) than those without (CT−TV−). Among *C. trachomatis*/*T. vaginalis*- coinfected participants, only 3 lacked detectable *M. hominis* or *Ca. M. girerdii*, limiting our ability to assess whether proinflammatory cytokines were further elevated when *T. vaginalis* harbored mycoplasmal endosymbionts. However, IL-8 was elevated in participants coinfected with *C. trachomatis* and endosymbiont-harboring *T. vaginalis* compared to those with *C. trachomatis* infection alone (Fig. 4C, P = 0.05).

**Figure 4.**
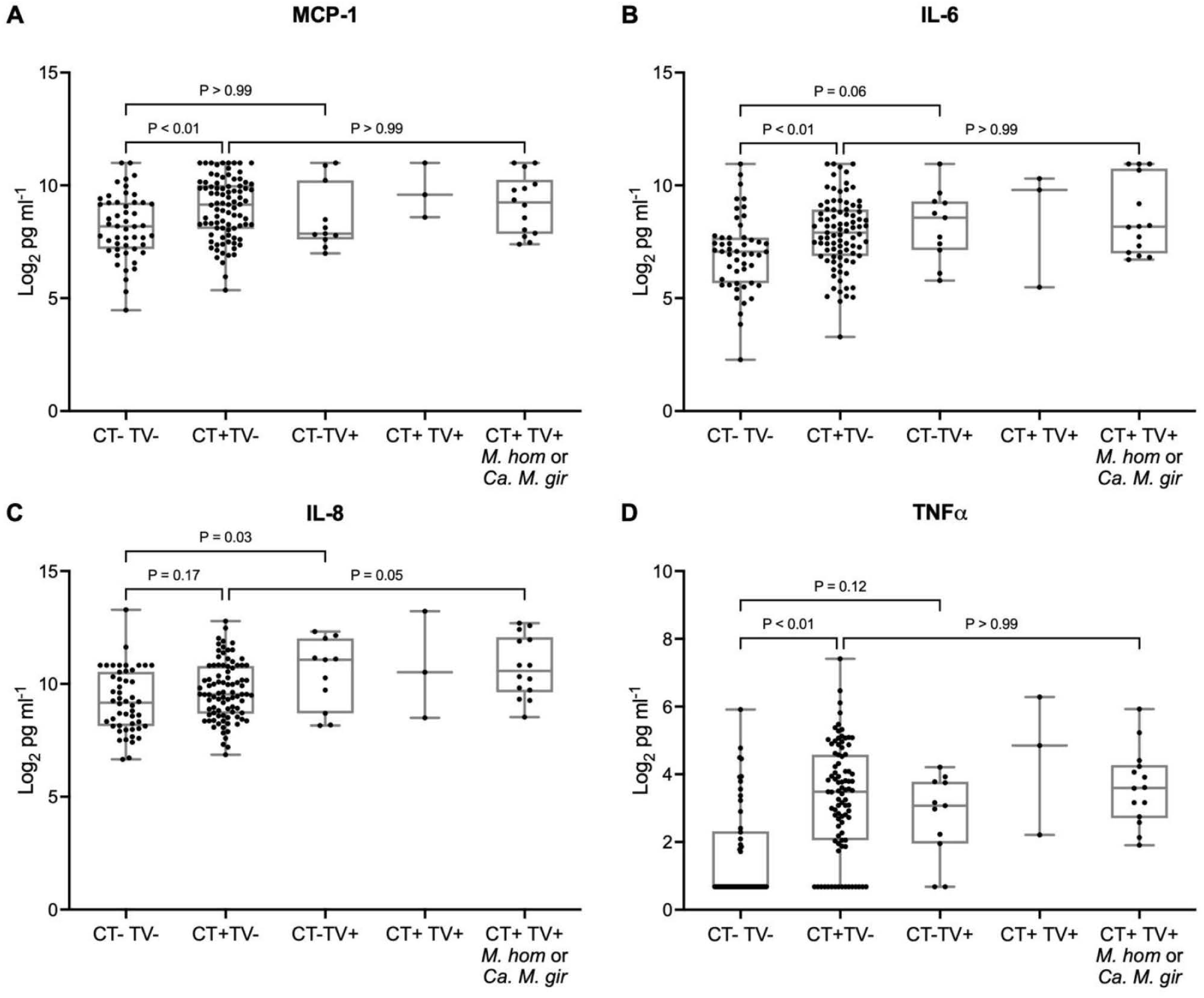
Cervical cytokine levels are elevated by chlamydial and trichomonad infection, alone and in combination, but are not further increased by mycoplasmal endosymbiont detection. Proinflammatory cytokines were quantified in cervical secretions from TRAC participants stratified by *C. trachomatis* (CT) and *T. vaginalis* (TV) infection status and endosymbiont ASV detection: uninfected (CT−TV−); CT monoinfected (CT+TV−); TV monoinfected (CT−TV+); CT/TV-coinfected without detectable *M. hominis* or *Ca. M. girerdii* ASVs (CT+TV+); and CT/TV-coinfected with detectable *M. hominis* and/or *Ca. M. girerdii* ASVs (CT+TV+ *M. hom/ Ca. M. gi*). Participants who tested positive for *N. gonorrhoeae* or *M. genitalium* at enrollment were excluded from this analysis. Levels of MCP-1/CCL2 (A), IL-6 (B), IL-8 (C), and TNFα (D) are shown for each group. Boxplots display distributions; Significance was determined by Kruskal-Wallis test followed by Dunn’s multiple comparisons test.

## Discussion

Pelvic inflammatory disease most often arises from bacterial infection of the uterus and fallopian tubes and can lead to serious reproductive sequelae including infertility, ectopic pregnancy, and chronic pelvic pain (35). Women with subclinical upper genital tract infection, “silent PID”, face similarly elevated risks (36). Coinfection with multiple STI pathogens is common (1, 37) but little is known about whether these pathogens interact synergistically or antagonistically to influence disease outcomes. Here, we investigated whether mycoplasmal endosymbionts of the sexually transmitted protozoan *T. vaginalis* modulated chlamydial upper genital tract infection in coinfected women. Using previously determined 16S rRNA gene ASV counts from vaginal swabs as estimates of endosymbiont abundance, alongside qPCR-based quantification of the protist, we found that *Ca. M. girerdii* and *M. hominis* ASVs trended toward more frequent detection, and higher abundance, in coinfected women without chlamydial spread to the endometrium, a relationship that may contribute to control of this obligate intracellular pathogen.

Although we lacked clinical specimens to confirm direct physical association between mycoplasmal bacteria and trichomonads in patients, we found that abundance of *M. hominis* and *Ca. M. girerdii* ASVs correlated significantly and positively with trichomonad DNA levels in individuals carrying both species and *M. hominis* alone, consistent with our hypothesis that these ASVs track *T. vaginalis* abundance. Evidence that mycoplasmal endosymbionts affect *T. vaginalis* abundance during natural infection is sparse, but consistent with other studies (13), we found that *Ca. M. girerdii* was detected almost exclusively in *T. vaginalis-*positive TRAC participants, consistent with its strict endosymbiotic nature. *M. hominis*, in contrast, was carried by 52% of participants who tested *T. vaginalis*-negative although at significantly lower mean abundance than in participants with trichomonad infection, suggesting that its free-living lifestyle is more nutritionally restricted and/or metabolically demanding. Abundance of the ASV identifying exclusively free-living *M. genitalium* was low or absent across the cohort, irrespective of *T. vaginalis-*coinfection, further underscoring the greater resource demands associated with a free-living existence for mollicutes. In vitro co-inoculation experiments have indicated that *M. hominis* and *Ca. M. girerdii* compete for host resources, affecting *T. vaginalis* growth and survival and potentially influencing overall parasite load (16). Amongst our cohort, many *T. vaginalis* infected participants carried both species, hinting at the potential complexity of trichomonad sub- populations occurring during natural infection.

Our prior analysis of factors contributing to chlamydial upper genital tract infection in TRAC found no evidence that *T. vaginalis* coinfection played a role (19), but that analysis relied on the binary outcome of *T. vaginalis* detection by molecular diagnostic test. We therefore re-examined whether *T. vaginalis* abundance, rather than mere presence, was associated with risk of chlamydial cervical or endometrial infection. We found no differences in chlamydial acquisition or upper tract spread based on *T. vaginalis* abundance overall. However, among coinfected participants, we observed a significant positive correlation between trichomonad abundance and chlamydial cervical burden, but only in women without coincident upper tract infection. Since chlamydial cervical burden is the strongest predictor of coincident endometrial infection in this cohort (20, 21) this raised the possibility that trichomonad populations in Endo− and Endo+ participants differ functionally in a way that is pathogenically relevant. We had examined mycoplasmal ASVs in relation to chlamydial endometrial infection for two reasons: they could be serving as surrogates for *T. vaginalis* in the original 16S rRNA gene study, and, as drivers of trichomonad metabolism, they have the potential to directly influence host abundance during infection. Nevertheless, endosymbiont-carrying *T. vaginalis* also display altered pathogenic potential: enhanced epithelial adherence in cell culture(16), synergistic upregulation of IL-8, IL- 1β, and TNF-α, and de novo induction of the Th17-polarizing cytokine IL-23 in human monocytic THP-1 cells (15, 17). We hypothesized that these proinflammatory mediators might help recruit immune cells to control a concurrent chlamydial infection and limit its spread to the upper genital tract, even in cases of high chlamydial burden. Consistent with this hypothesis, IL-8 was elevated in *C. trachomatis*/*T. vaginalis*-coinfected participants with detectable mycoplasmal endosymbionts compared to participants with *C. trachomatis* infection alone (P = 0.05). In contrast, MCP-1/CCL2, IL-6, and TNFα were not further elevated by endosymbiont carriage beyond what was already seen with *T. vaginalis* infection alone. IL-8 (CXCL8) is a potent chemoattractant for neutrophils, and IFN-γ-stimulated neutrophils have been shown to kill antibody-opsonized (38). Thus, IL-8 mediated neutrophil influx could contribute to reduced endometrial chlamydial spread. Unfortunately, we could not directly compare cytokine levels between endosymbiont-positive and endosymbiont-negative coinfected women because only 3 coinfected participants lacked detectable *M. hominis* or *Ca. M. girerdii*.

Our ASV data independently support a role for endosymbiont carriage in limiting chlamydial spread. *Ca. M. girerdii* ASV was detected more frequently in Endo− than Endo+ women (p = 0.04) among women with high chlamydial burden (≥ 10^4^ GE), a difference not observed among women with low chlamydial burden (p = 1.0). We also observed trends for presence and abundance of both endosymbionts toward association with absence of chlamydial upper tract infection, along with a significant positive association between *M. hominis* ASVs and trichomonad abundance among Endo− participants. Notably, *M. hominis* ASVs were not selected as classifiers in our original 16S rRNA gene study predicting ascended chlamydial infection but when considered in the context of *T. vaginalis*— where *M. hominis* has the potential to live endosymbiotically — we detected a strong, statistically significant, correlation of *M. hominis* ASV with trichomonad abundance in women without endometrial infection. Together, these findings point to a shared functional feature of trichomonads harboring endosymbionts that may modulate chlamydial infection dynamics at the cervix through increased induction of a neutrophil chemoattractant, as well as other mechanisms — consistent with a broader literature implicating bacterial co-pathogens and cervicovaginal microbiome members in synergistic or antagonistic effects on chlamydial pathogenesis (reviewed in Hand et al. (39)). In the case of *N. gonorrhoeae*, evidence suggests that despite competing with chlamydiae for iron, it suppresses developing adaptive immunity by promoting antigen-presenting cell death and restricting T-cell proliferation (40, 41), synergizing with chlamydial expression of Chlamydial Protease Activity Factor (CPAF), a potent suppressor of neutrophil function (42), to the benefit of both pathogens. For coinfecting

*T. vaginalis*, regardless of endosymbiont status, abundant local secretion of CPAF would likely benefit the protist since neutrophils are heavily involved in controlling the parasite through trogocytosis (43) and other mechanisms (44).

Our study had several limitations. First, we lacked information on the presence or abundance of *T. vaginalis*-specific double-stranded RNA viruses (TVVs) in study participants (45, 46). Four TVV species, members of the Totiviridae, are known to enhance proinflammatory signaling via TLR3 when virions are released from stressed or dying protozoa (47) and may have served as potential confounders of cytokine secretion we could not account for. Second, as noted above, we lacked appropriate biospecimens to directly validate our inferences about an endosymbiotic versus free-living lifestyle for *M. hominis* in trichomonads from Endo− women. Competition between endosymbiotic *M. hominis* and *Ca. M. girerdii* during dual infection of *T. vaginalis*, which can affect parasite metabolism and cytoadhesion (16), may also have modulated outcomes in these participants. Finally, statistical power was limited: even among participants with *T. vaginalis* monoinfection, the vast majority (9/10) carried endosymbiont ASVs, leaving too few individuals to meaningfully stratify by subgroup.

Future cohort studies with enrichment for relevant coinfection populations could help address these gaps. We are actively investigating the cervicovaginal microbiome of TRAC2 participants (48), which may offer an opportunity both to validate our findings and to enable future transcriptional or cell-profiling studies of *T. vaginalis* coinfection subgroups.

## Acknowledgements

We thank the women who agreed to participate in this study; Ingrid Macio, Melinda Petrina, Carol Priest, Abi Jett, and Lorna Rabe for their efforts in the clinic and the microbiology laboratory; and the staff at the Allegheny County Health Department STD Clinic, for their efforts. We also acknowledge the contribution of Daniel Fitzgerald to optimization of qPCR assay. The authors also acknowledge Dr Andrew Macintyre, Manager and Lead Investigator in the Immunology Unit of the Duke Regional Biocontainment Laboratory for cytokine assays.

## Authors’ contributions

TD, SJ, XP, KSY and CMO designed the study. TT, CC, and CMO performed the experiments. KSY and NB performed the data curation. XS, NB, and XZ performed the data analysis. KSY, TD, and CMO wrote the manuscript. All authors reviewed the manuscript.

---

**Figure S1.**
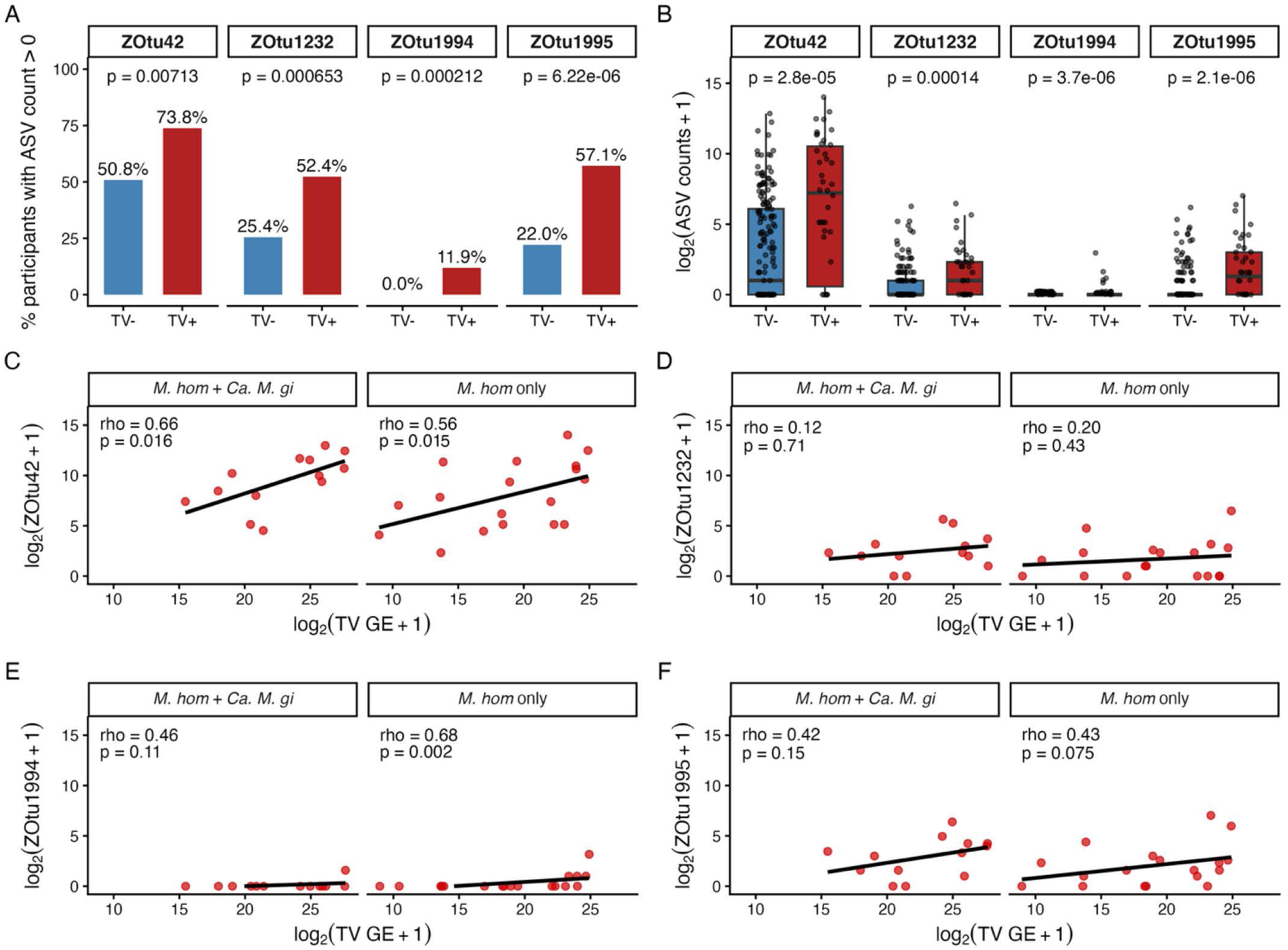
Association between individual amplicon sequence variants (ASVs) assigned to *M. hominis* and *Trichomonas vaginalis* (TV) infection status and abundance among TRAC participants. (A) Percentage of participants with detectable ASV counts (>0) for each M. hominis ZOtu, comparing TV− and TV+ groups. Bars show proportions; P-values were obtained by chi-square or Fisher’s exact test. (B) Abundance of each ZOtu (log₂[count + 1]), stratified by TV status, with Wilcoxon rank-sum P-values. (C–F) Association between *T. vaginalis* burden (log₂[TV genome equivalents + 1]) and ZOtu-specific ASV counts among TV-positive participants, stratified into participants co-colonized with *M. hominis* and *Ca. M. girerdii* (*M. hom*/ *Ca. M. gi*) versus those carrying *M. hominis* alone (*M. hom*-only). Each panel shows individual participant values, a fitted linear regression line, and the Spearman correlation coefficient (ρ) with P-value. Panels correspond to (C) ZOtu42, (D) ZOtu1232, (E) ZOtu1994, and (F) ZOtu1995.

